# Structure of Native Chromatin Fibres Revealed by Cryo-ET *in situ*

**DOI:** 10.1101/2023.09.03.556082

**Authors:** Zhen Hou, Frank Nightingale, Yanan Zhu, Craig MacGregor-Chatwin, Peijun Zhang

## Abstract

The structure of chromatin plays pivotal roles in regulating gene transcription, DNA replication and repair, and chromosome segregation. This structure, however, remains elusive. Using cryo-FIB and cryo-ET, we delineated the 3D architecture of native chromatin fibres in intact interphase human T-lymphoblasts and determined the *in-situ* structures of nucleosomes in different conformations. These chromatin fibres are not structured as uniform 30 nm one-start or two-start filaments but are composed of relaxed, variable zigzag organizations of nucleosomes connected by straight linker DNA. Nucleosomes with little H1 and linker DNA density were distributed randomly without any spatial preference. This work sets a precedent for future high-resolution investigations on native chromatin structures *in-situ* at both a single-nucleosome level and a population level under many different cellular conditions in health and disease.

## Introduction

In eukaryotes, chromatin is a highly dynamic nucleoprotein complex that not only stores genetic information, but also participates in gene expression, DNA replication, and DNA repair. Chromatin can undergo drastic changes in structure and composition during the cell cycle and in response to various environmental and cellular signals^1-5^. The building block of chromatin is the nucleosome, consisting of a 1.65-turn wrapped 147-base-pair DNA string and an octamer of histones, H2A, H2B, H3, and H4 histone dimers^6-9^. However, to accommodate a two-meter-long DNA string into a mammalian nucleus of around 10-μm dimeter^10^, nucleosomes must be further packed into higher-order chromatin structures.

There have been extensive studies on chromatin fibres and nucleosome compaction. Purified chromatin at low ionic strength can be seen as connected nucleosomes, presenting a 10-nm beads-on-a-string structure which further coils into 30-nm chromatin fibres in the presence of linker histone H1 under moderate ionic conditions^11-15^. There have been several models proposed for the 30-nm chromatin fibre based on *in vitro* studies, the two most prominent being the zigzag and solenoid models^15-25^. The zigzag model, also known as the two-start fibre model (including both the “helical ribbon” and “twisted crossed-linker” models) suggests nucleosomes zigzag back and forth with relatively straight DNA linkers^18-20, 22, 23^. The solenoid model, also known as the one-start fibre model, suggests a helical structure generated by nucleosomes stacking linearly along the helical axis, where the linker DNA is bent connecting adjacent nucleosomes^17, 21, 25^. Computer simulations have suggested the coexistence of both models as well as flexible disordered models in the nucleus^26-29^. However, the existence of 30-nm chromatin fibres in the native nucleus has long been debated over the past decades, as no such defined fibres have been observed in intact cells under native conditions^30-35^. The structure of native chromatin fibres thus remains elusive. Here we investigated how nucleosomes are organized into chromatin fibres within the intact frozen-hydrated T-lymphoblast CEM cell nucleus *in-situ* using cryo-focused ion beam (cryo-FIB) and cryo-electron tomography (cryo-ET).

## Results

### Chromatin fibres revealed in the intact nucleus

To visualise the chromatin, we generated very thin CEM cell lamellae containing the interphase nucleus by automated cryo-FIB milling, with the thinnest lamellae about 80-90 nm thick. The reconstructed tomograms clearly resolve chromatin fibres and individual nucleosomes in the heterochromatin region close to the nuclear envelope (Fig. 1a-d, Supplementary Movie 1). The width of these fibres is variable, ranging from 20 to 50 nm (Fig.1a, c, d, Fig. 2c). An array of nucleosomes was seen to display a similar structure as the *in vitro* reconstituted two-start zigzag complex (Fig. 1c, circle)^23^ and in vitreous sections of isolated chicken erythrocyte nucleus^36^. Moreover, individual nucleosomes can be identified flanking a DNA spine (Fig. 1d, yellow oval). Nearly naked, nucleosome-free DNA was also observed (Fig. 1d, yellow arrowhead), which continues as a chromatin fibre. These direct observations from individual nucleosomes, together with the distributions of pairwise distances and angles between adjacent nucleosomes from a large population (Figure 1F), indicate that the native chromatin fibres appear highly variable and flexible, and no uniform 30 nm fibre structure was observed.

**Figure 1.**
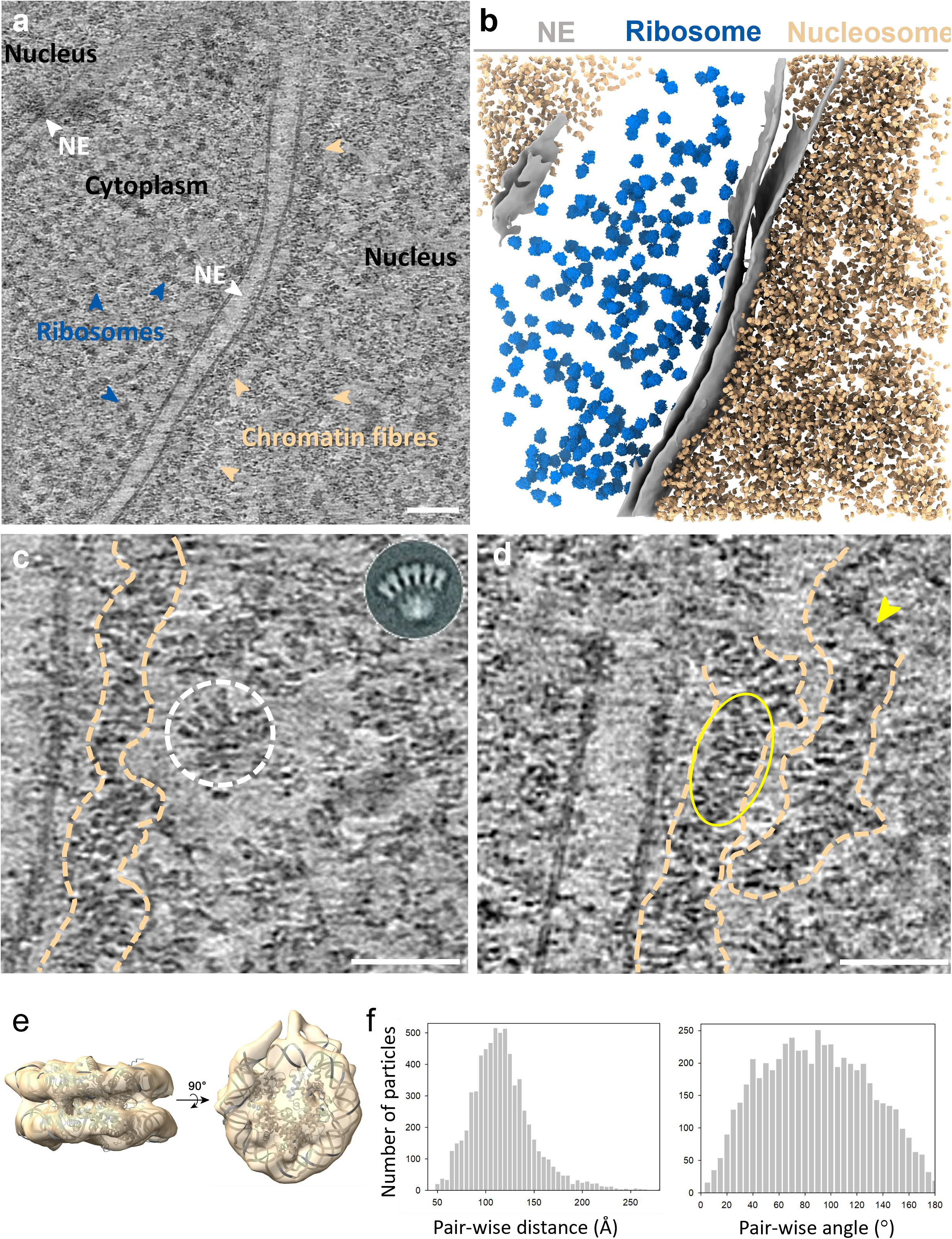
Cryo-ET of native chromatin fibres and subtomogram average of nucleosomes. **a)** A representative tomographic slice of the CEM cell (from *n* = 5). Scale bar = 100 nm. The tomogram is reconstructed with SIRT-like filtering in IMOD 4.11. The nucleus, chromatin fibres, nuclear envelope (NE), cytoplasm, and ribosomes are labelled accordingly. **b)** The segmented volume of the tomogram in **(a)**. NE, ribosomes and nucleosomes are coloured grey, blue and gold, respectively. **c-d)** Representative tomographic slices of chromatin fibres in the nucleus (from *n* = 5). Prominent fibre structures are indicated by dash outlines. The white dashed circle showcases a nucleosome array similar to the two-start zigzag structure of the reconstituted nucleosome complex (inset) (ref. ^23^). The yellow arrowhead points to naked DNA and the yellow oval shows nucleosomes franking a DNA spine. Scale bars = 50 nm. **e)** *In situ* structure of native nucleosomes (from n = 6,790, n of tomograms = 5), fitted with a crystal structure of core nucleosome (PDB 6ESF), shown in two orthogonal views. **f)** Distributions of pair-wise distances and angles between the nearest nucleosome neighbours (from *n* = 6,790, *n* of tomograms = 5).

**Figure 2.**
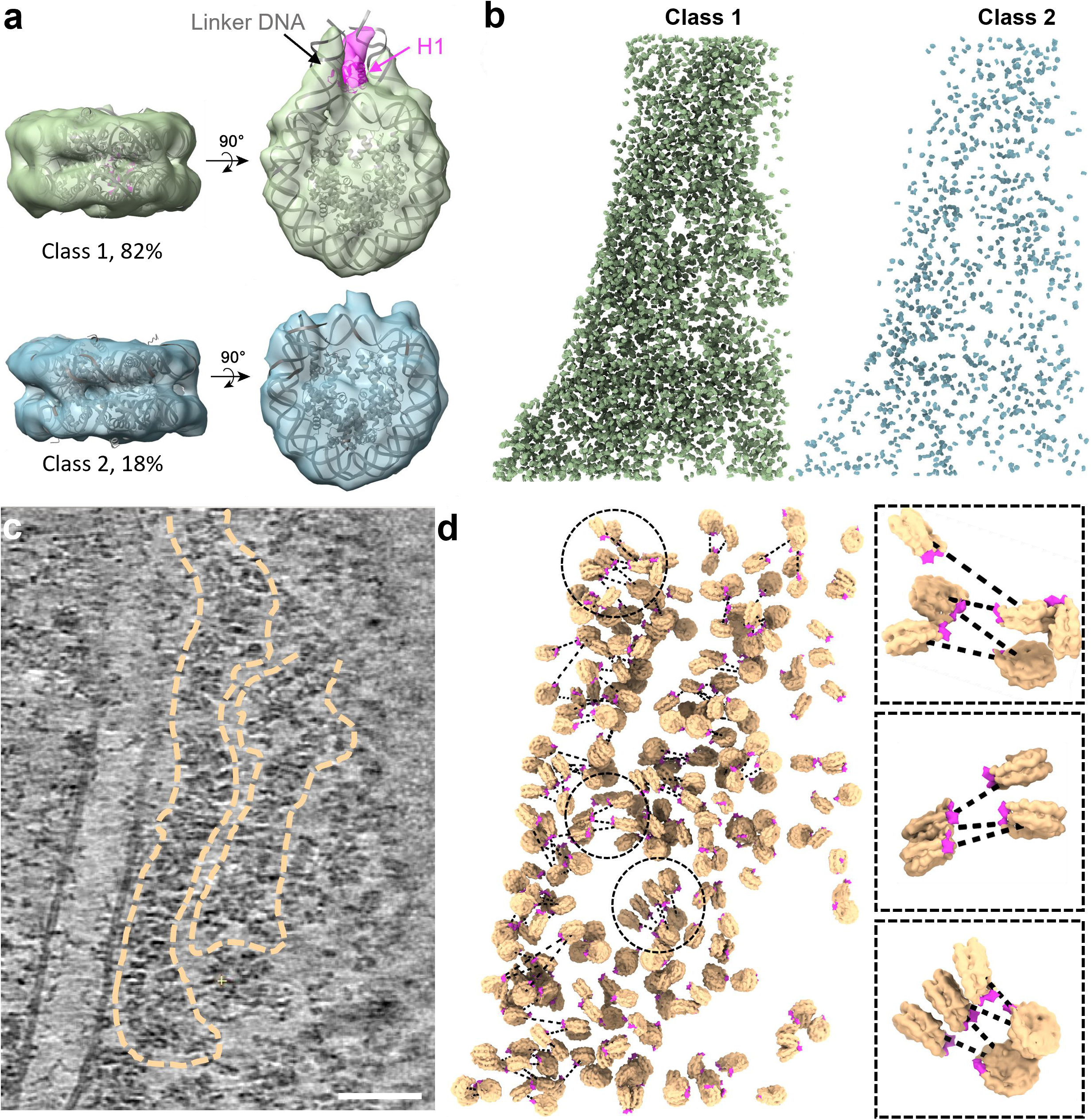
3D Organization of nucleosomes in native chromatin fibres. **a)** Two distinct classes of native nucleosomes. Class 1 (top) (82% of the total population, *n* = 5,578) is fitted with the nucleosome crystal structure PDB 7DBP, with H1 density coloured in magenta. Class 2 (bottom) (18% of the total population, *n* = 1,212) is fitted with the core nucleosome crystal structure PDB 6ESF. **b)** Mapping the back of individual nucleosomes from Class 1 (left) and Class 2 (right) into the representative tomogram according to their coordinates and orientations. **c)** A rotated tomographic slice (Y axis: -16°, from Fig. 1d) depicting two clear chromatin fibres (dashed outlines). Scale bar = 50 nm. **d)** Mapping back of individual nucleosomes in chromatin fibres in (**c**). The H1 density is coloured magenta, indicating the side of entry and exit of the linker DNA, from which the DNA path (dashed lines) is predicted. Three representative sub-regions (circled) are enlarged on the right panel.

### *In-situ* structures of nucleosomes in the native nucleus

Template matching using a featureless nucleosome model (EMD-26339 with a 40 Å low-pass filter)^37^ resulted in 10,000 particles from 5 tomograms of thinnest lamellae. After the removal of false positives by classification and manual inspection, the remaining 6,790 nucleosome particles were iteratively aligned and yielded a subtomogram average of the *in-situ* nucleosome at 12.0 Å resolution (Fig. 1e, Supplementary Fig. 1a, b, Supplementary Table 1, Supplementary Movie 2). The DNA dyads are clearly differentiated along with partial densities of the linker DNA and linker histone H1 (Fig. 1e). Further 3D classification of nucleosome particles resulted in two major distinct classes: class 1 (82%) shows a clear linker DNA density with partially resolved H1 globular domain (Fig. 2a, top), whereas class 2 shows little linker DNA and H1 density, suggesting these nucleosomes are likely more dynamic or flexible and perhaps have also lost H1 (Fig. 2a, bottom, Supplementary Fig.1c, d). To test whether these two distinct populations of nucleosomes form special domains or prefer certain localization, we mapped each nucleosome from both classes back to the original tomogram. Intriguingly, the nucleosome particles from each class are distributed rather randomly without any spatial preference (Fig. 2b).

### 3D organization of nucleosomes in native chromatin fibres

The high quality of tomograms allowed for the investigation of the architecture of the native chromatin fibre. As the individual chromatin fibres can be delineated along with the nucleosomes in the original tomograms (Fig. 1c, d, Fig. 2c, marked with dashed lines, Supplementary Movie 3), we placed all identified nucleosomes back into the fibre tomogram according to their refined positions and orientations (Fig. 2d). The nucleosome model used for placement specifies the position of H1 and linker DNA (Fig. 2d, magenta), and thus allows prediction of the DNA path and connection of adjacent nucleosomes (Fig. 2d, dashed black lines).

Within the fibres, the median distance between adjacent nucleosomes was measured at ∼120 Å, which is significantly larger than the expected distance (60-90 Å) calculated from the compact fibre models^17-20, 23, 25^ (Supplementary Fig. 2a). Moreover, when we divided nucleosomes into three subgroups of neighbouring distances, 60-80 Å, 80-100 Å, 100-120 Å, none of the nucleosome subpopulations showed a compact uniform fibre structure (Supplementary Fig. 2b). Rather, all of them exhibited a flexible zigzag configuration. Consistently, the wide range of pairwise angles (Fig. 1f), and the distribution of nucleosome subpopulations with different ranges of angles (Supplementary Fig. 2c), further support that the chromatin fibres are largely made of non-uniform, flexible zigzag-arranged nucleosomes. The calculated DNA concentration in our tomogram is about 15 mg/ml, which is consistent with the concentration of DNA within the nucleus (∼10 mg/ml), suggesting the model is able to pack the entire genome into the nucleus. While the analysis of the overall nucleosome population does not suggest a uniform compact chromatin fibre, there are a few instances of short-ranged, partial compact nucleosomes (Fig. 1c, 2d), including di-nucleosomes joined via interface 1 or interface 2^23^ (Supplementary Fig. 2d1-3), tetra-and poly-nucleosomes (Fig. 2d, Supplementary Fig. 2d4-5).

## Discussion

The chromatin fibre and its structural model have been investigated and debated for many years. Using state-of-the-art cryo-FIB and cryo-ET, we revealed the elusive architecture of native chromatin fibres in intact cells without the drastic manipulation and harsh treatment of samples of previous studies^38, 39^, and determined the *in-situ* structure of nucleosomes at unprecedented resolution of 12 Å, which is likely limited by the intrinsic dynamics and heterogeneity of native nucleosomes. The capacity to resolve linker DNA and H1, in partial, was critical to elucidating how nucleosomes are connected, and thus the structure of continuous chromatin fibres, which has not been possible in previous efforts^31,32,35^. Our data show that most nucleosomes are connected by straight linker DNA, forming a flexible, relaxed zigzag pattern, substantiated by both direct visualizations of individual nucleosomes and large-scale nucleosome population analysis. The architecture of the chromatin fibre determines DNA accessibility for transcription and other template-directed biological processes. Non-rigid chromatin fibre is likely beneficial for the effective tuning of the genome in response to varying protein expression and cellular stresses.

Chromatin per se constantly transforms during the entire cell cycle in response to various cell signalings^1, 4, 5, 40^. While the structure of chromatin and chromatin fibres vary with cell types, cell states, nuclear positions, and signal perturbations, the fibre model we have postulated suggests a general mechanism for the compaction of the genetic material. The revelation of chromatin fibre structure in the interphase mammalian nucleus opens a new avenue for future high-resolution *in-situ* investigations of various native chromatin structures and their relevance to gene expression, the cell cycle, and stress responses.

## Materials and Methods

### Cell culture and vitrification

CEM CD4+ T-cells (catalogue ARP-117, HIV reagents program) were cultured in DMEM (Gibco) supplemented with 10% FBS, 2mM L-glutamine (Gibco) and 1% MEM nonessential amino acids (Gibco), at 37°C and 5% CO_2_. CEM cells at 3x10^6^ cells/ml after 10 passages were pelleted at 200 g for 5min at 20°C and resuspended in PBS mixed with 10% glycerol. An aliquot of 3 ul cell suspension was applied to the glow-discharged holey carbon-coated copper (R 2/1, 200 mesh) (Quantifoil) and blotted for 9 seconds by Leica GP2 (Leica Microsystems), followed by plunge freezing in liquid ethane.

### Cryo-FIB milling

Vitrified cells were further processed by cryo-FIB milling for the preparation of lamellae. A dual-beam microscope FIB/SEM Aquilos 2 (Thermo Fisher Scientific) equipped with a cryotransfer system (Thermo Fisher Scientific) and rotatable cryo-stage cooled at -191°C by an open nitrogen circuit was used to carry out the thinning. Prior to the milling, the grids were mounted on the shuttle and transferred onto the cryo-stage, followed by the coating with an organometallic Platinum layer using the GIS system (Thermo Fisher Scientific) for 5-6 seconds. Then, cells positioned approximately in the centres of grid squares were selected for thinning. The thinning was conducted by the automated milling software AutoTEM 5 (Thermo Fisher Scientific) in a stepwise manner from current 0.5 μA to 30 pA at 30 kV, and the final thickness of lamellae was set to 120 nm.

### Cryo-ET data collection

Cellular lamellae were transferred to FEI Titan Krios G2 (Thermo Fisher Scientific) electron microscope operated at 300 kV and equipped with a Gatan BioQuantum energy filter and post-GIF K3 detector (Gatan, Pleasanton, CA). A 100 μm objective aperture was inserted. Areas that include nuclei were selected for the data acquisition. Tilt series were recorded using Tomography 5 software (Thermo Fisher Scientific) with a nominal magnification of 42k and a physical pixel size of 2.18 Å/pixel. All tilt series were collected with a zero-loss imaging filter with a 20eV-wide slit. The defocus value was set from -3.5 to -5 μm. The pre-tilt of the lamellae was determined at + 9°, and a dose-symmetric scheme was applied for all tilt series, ranging from -45° to +63° with an increment of 3°. A total of 37 projection images with 10 movie frames each were collected for each tilt series and the dose rate was set at 1.5 e/Å^2^/s with an exposure time of 2 seconds, resulting in a total dose of 111 e/Å^2^. The correlated double sampling (CDS) in super-resolution mode was applied and frames were saved in LZW compressed tif format with no gain normalization. A total of 26 tilt series were collected from 22 lamellae.

### Alignment of tilt series and tomogram reconstruction

The frames of each tilt series were corrected for beam-induced motion using MotionCor2^41^. The gain correction was performed in parallel with the motion correction run by a home-brewed script. New stacks were generated and aligned using IMOD^42^ version 4.11 by patch tracking, and tomograms were reconstructed at bin6 with a pixel size of 13.08 Å/pixel. For visualisation and segmentation, reconstructed tomograms were corrected for missing wedge and denoised by IsoNet^43^ version 0.2, applying default parameters.

### Template matching

To localise individual nucleosomes in the tomogram, template matching was carried out using emClarity^44^ version 1.5.0.2. To supress the template-induced bias, a featureless nucleosome template was generated by low-pass filtering the published structure EMD26339^37^ to 40 Å. A total number of 5 tomograms with low residual errors in the alignment were selected for template matching. About 10000 particles in total were extracted. In parallel, ribosomes were picked using the low-pass-filtered structure EMD-16196^45^, and 200 particles were extracted from each tomogram for the segmentation.

### Subtomogram averaging

Prior to aligning the particles, CTF correction was performed for each tomogram by emClarity^44^ version 1.5.3.10, and particles were checked by overlaying the reconstructed tomograms with corresponding picked particles in Chimera. Particles that lay outside the nucleus were then removed. The remaining particles were first aligned at bin6 and bin5 by emClarity version 1.5.3.10, followed by iterative reconstructions and alignments at lower binning (2-4) in RELION^46^ version 4.0. To further clean up the particles, 3D classification was conducted at bin 6 in RELION^46^ version 4.0, and 6,790 particles remained after the cleaning. The final resolution of the nucleosome was determined at 12.0 Å (0.143 cut-off). Using that structure as the reference, another round of 3D classification in RELION^46^ was performed at bin6 with the number of classes set as 10. Class 1, 3, 4, 5, 6, 7, 8, and 9 showed prominent linker DNA densities while classes 2 and 10 did not, thus particles from class 1, 3, 4, 5, 6, 7, 8, 9 were combined as one class (Class 1: 5,578 particles), particles from class 2 and 10 were combined as the other class (Class 2: 1,212 particles) (Supplementary Figure 1C). These two classes were then aligned iteratively, and the final resolution of class 1 was determined at 12.5 Å (gold-standard 0.143 cut-off) (Supplementary Figure 1B, blue line) while class 2 was resolved at 15 Å (gold-standard 0.143 cut-off). Map fitting was performed in ChimeraX^47^, and the PDB 7DBP^48^ and PDB 6ESF^49^ were compared with class 1 and class 2, respectively.

### Analysis of nucleosome population

The distance between adjacent nucleosomes was calculated according to the coordinates of their centres after the refinement. Paired nucleosomes with a centre-to-centre distance shorter than 60 Å were regarded as duplicates and removed. The angle between non-duplicate neighbouring nucleosomes was calculated using an in-house-developed script (https://github.com/fnight128/MagpiEM).

### Segmentation and visualization

The segmentation was performed on IsoNet^43^-processed tomograms at bin6. The initial membrane detection and segmentation were done by TomoSegMemTV^50^, and the successive rendering was accomplished by ChimeraX^47^. Nucleosomes were mapped back according to their refined coordinates and orientations, using the subtomogram averaged nucleosome structure as the model. Ribosomes were mapped back to the tomogram as well, using a lowpass-filtered structure of EMD-16196^45^ as the model. Positions and orientations of ribosomes were based on the outputs from template matching.

## Data availability

The *in-situ* nucleosome structures have been deposited in EMDB under the accession codes EMD-16978, EMD-16979 and EMD-16980 for all nucleosomes, Class 1 nucleosomes, and Class 2 nucleosomes, respectively.

## Code availability

The scripts used in this study and relevant codes are deposited in GitHub: https://github.com/fnight128/MagpiEM.

## Acknowledgement

We thank Stanley Fronik for help in sample preparation, we thank Dr. Karen Davies for her support in data collection, we thank Dr. Nathan Hardenbrook for help with data analysis, and also acknowledge Dr. Long Chen for suggestions in statistical analyses. We acknowledge The Oxford Particle Imaging Centre (OPIC) for access to the cryo-FIB/SEM instrument (Aquilos 2) and Diamond Light Source for access and support of the cryo-EM facilities at the U.K. national eBIC (proposal NT29812), funded by the Wellcome Trust, MRC, and BBSRC. This research was supported by the UK Wellcome Trust Investigator Award (206422/Z/17/Z), the National Institutes of Health grants AI150481, the ERC AdG grant (101021133), and the Wellcome Trust Core Award Grant Number 203141/Z/16/Z with additional support from the NIHR Oxford BRC.

## Author contributions

P.Z. conceived the research. Z.H. and P.Z. designed the experiments. Z.H. and C.M. prepared the lamellae. Z.H. and Y.Z collected data. Z.H. carried out tomography reconstruction and subtomogram averaging and classification. Z.H., F.N. and P.Z. analysed the data. Z.H. and P.Z. wrote the manuscript with help from all co-authors.

