## supplementary figures for "Structure of Native Chromatin Fibres Revealed by Cryo-ET *in situ*"

### Supplementary information

1. Supplementary Table 1
2. Supplementary Figures 1-2
3. Supplementary Movies 1-3

**Supplementary Table 1 |** CryoET data collection and structure determination of native nucleosomes

| **Sample** | T-lymphoblast CEM CD4+ cell lamellae |
|---|---|
| Microscope | FEI Titan Krios |
| Voltage (keV) | 300 |
| Detector | Gatan Quantum K3 Direct Electron Detector |
| Energy-filter | Yes |
| Slit width (eV) | 20 |
| Super-resolution mode | No |
| Physical pixel size (Å/pixel) | 2.18 |
| Defocus range (µm) | -3.5 to 5, increment 0.3 |
| Acquisition scheme | Dose-Symmetric, -45º to 63º, 3º step, group 3 |
| Total dose (electrons/Å$^2$) | 111 |
| Number of frames | 10 |
| Number of cells | 5 |
| Number of tomograms | 5 |
| **CryoET and subtomogram averaging** | |
| Number of subtomograms | 6790 |
| Resolution at 0.143 FSC cut-off (Å) | 12.0 |
| Data deposited | EMD-16978, EMD-16979, EMD-16980 |

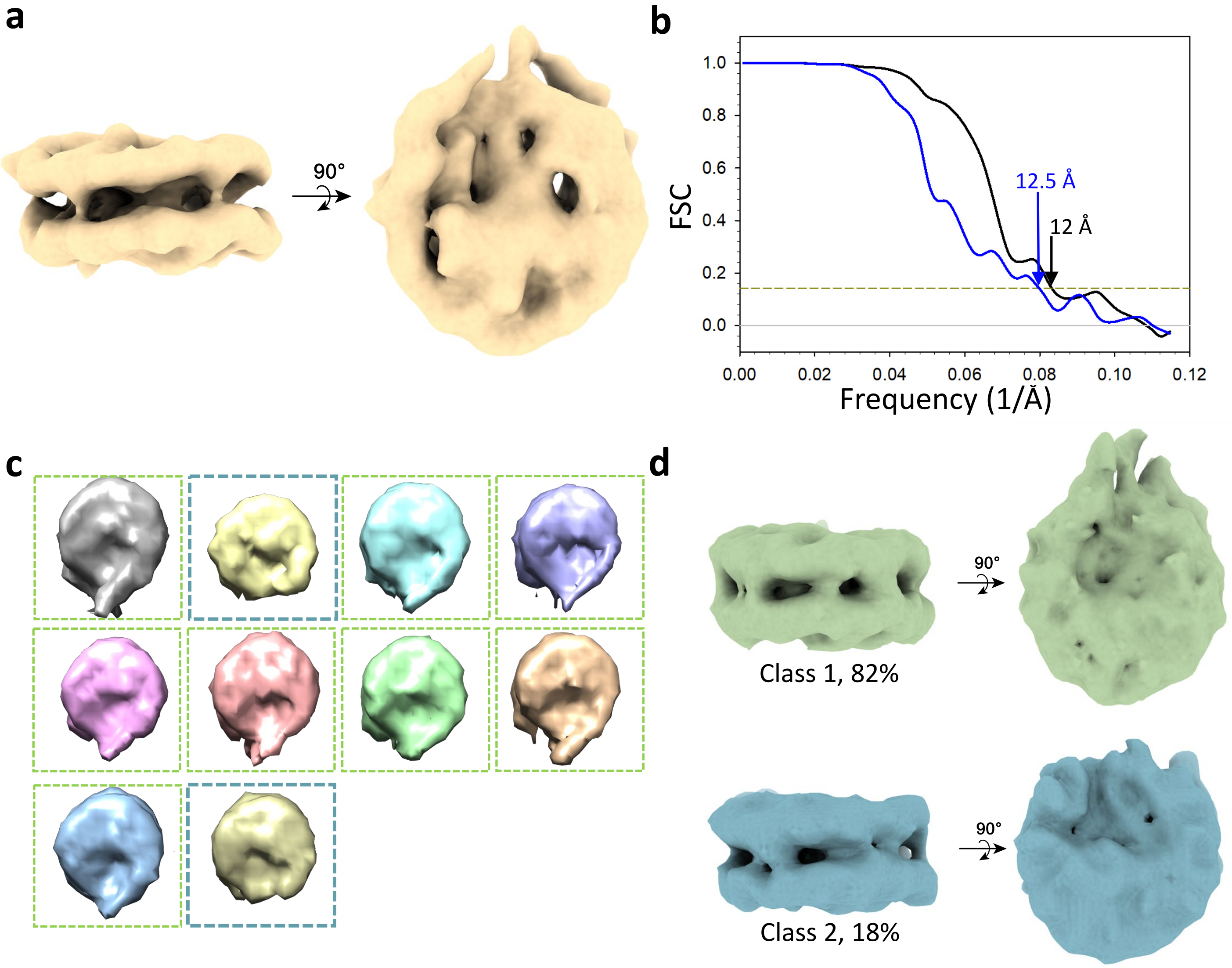


**Supplementary Figure 1 | Subtomogram averaging and classification of native nucleosomes. a)** A subtomogram average of native nucleosomes *in situ* (from *n* = 6,790, *n* of tomograms = 5). Two orthogonal views are shown. **b)** Gold-standard Fourier shell correlation (FSC) curves of subtomogram averaged maps from all nucleosomes (black line) and from Class 1 (blue line). **c)** Classification of 6,790 native nucleosome particles. Nucleosome particles from classes framed by dashed green lines showing prominent linker DNA density and partial H1 density are combined into Class 1, whereas nucleosome particles from classes framed by light blue lines corresponding to the canonical core nucleosome structure are combined into Class 2. **d)** Subtomogram averages of Class 1 nucleosomes (from *n* = 5,578) (top) and Class 2 native nucleosome (from *n* = 1,212) (bottom), shown in two views.

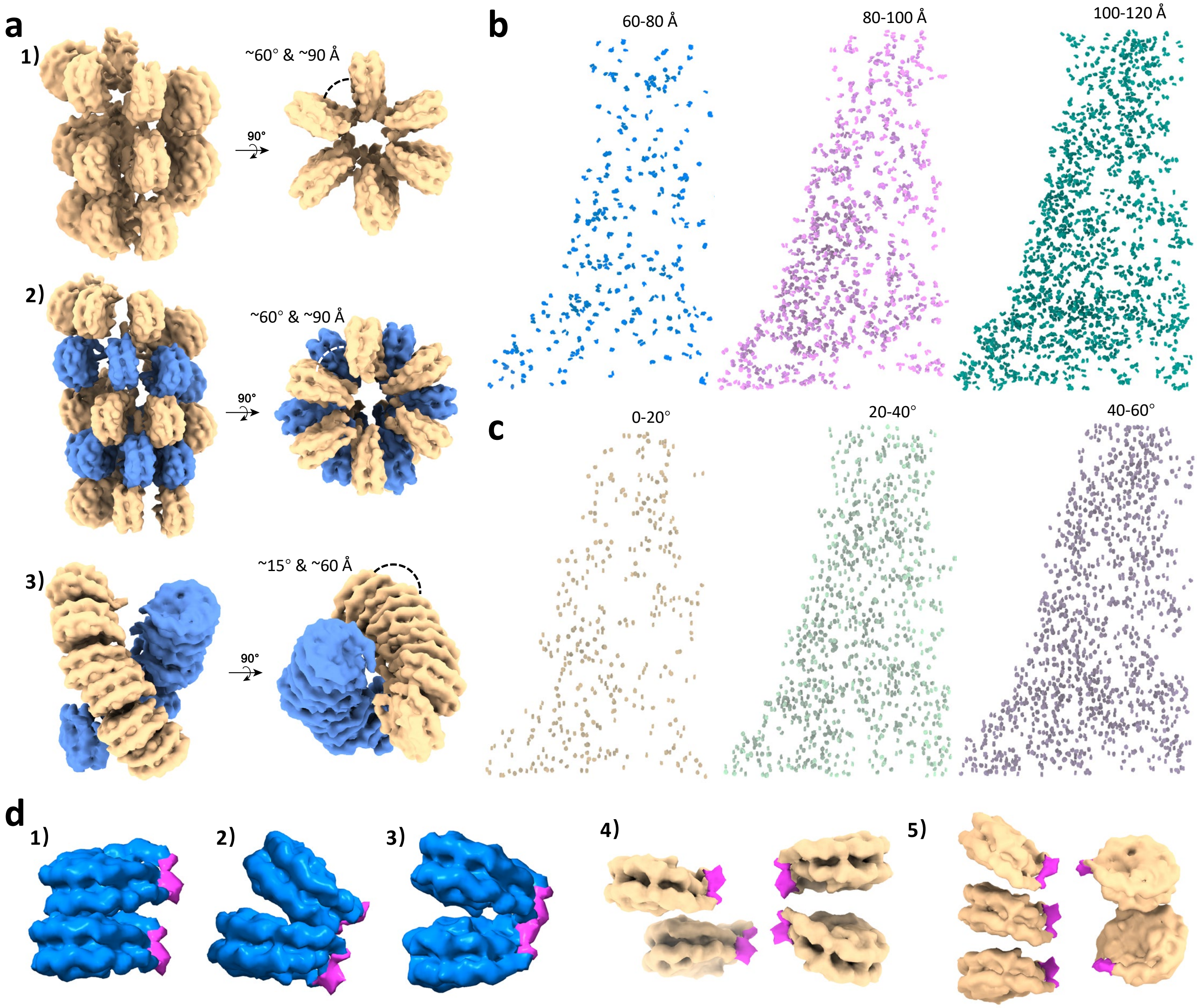


**Supplementary Figure 2 | Chromatin models and mapping of different subpopulations of the native nucleosome. a)** Uniformed chromatin fibre models constructed based on previous works: 1) Ideal one-start solenoid model, 2) Ideal two-start twisted crossed-linker zigzag model, 3) EM-based compact two-start twisted zigzag model. The distance and angle between the neighboring nucleosomes are indicated. **b)** Mapping back of three subpopulations of nucleosomes based on the nearest neighbor distance: 60-80 Å (left), 80-100 Å (middle), 100-120 Å (right). **c)** Mapping back of three subpopulations of nucleosomes based on the angle between the nearest neighbours: 0-20° (left), 20-40° (middle), 40-60° (right). **d)** Examples of partial compact di-nucleosomes with type I (1) and type II interactions (2-3) (ref. [23]), tetra- (4) and poly-nucleosomes (5) from the subpopulation of 80-100 Å in (**b**). Partial densities of linker DNA and H1 are colored magenta.

**Supplementary Movie 1 | A representative reconstructed tomogram with the segmentation of nuclear envelope and mapping of ribosomes and nucleosomes.** The tomogram is reconstructed with Sirt-like filtering in IMOD 4.11 and sliced along the Z axis back and forth through slice 1 to slice 269. The segmented volume is overlapped with the tomogram in ChimeraX, NE, ribosomes and nucleosomes are coloured grey, blue and gold, respectively. An enlarged view of the nucleosome is introduced at the end.

**Supplementary Movie 2 | *In situ* structure of the native nucleosome fitted with the crystal model.** The transparency of the *in situ* structure is adjusted to 50% and the crystal model (PDB 6ESF) is coloured grey. The spinning is 360° and conducted in ChimeraX.

**Supplementary Movie 3 | Representative tomographic slices of native chromatin fibres.** The reconstructed tomogram is fixed at -16° of the Y axis and rotated back and forth along the X axis from -15° to +15° in IMOD 4.11.
